# Noncontractile tissue forces mask muscle fiber forces underlying muscle spindle Ia afferent firing rates in stretch of relaxed rat muscle

**DOI:** 10.1101/470302

**Authors:** Kyle P. Blum, Paul Nardelli, Timothy C. Cope, Lena H. Ting

## Abstract

Stretches of relaxed cat and rat muscle elicit similar history-dependent muscle spindle Ia firing rates that resemble history-dependent forces seen in single activated muscle fibers (Nichols and Cope, 2004). During stretch of relaxed cat muscle, whole musculotendon forces exhibit history-dependence that mirror history-dependent muscle spindle firing rates, where both muscle force and muscle spindle firing rates are elevated in the first stretch in a series of stretch-shorten cycles (Blum et al., 2017). By contrast, rat musculotendon are only mildly history-dependent and do not mirror history-dependent muscle spindle firing rates in the same way (Haftel et al., 2004). We hypothesized that history-dependent muscle spindle firing rates elicited in stretch of relaxed rat muscle would mirror history-dependent muscle fiber forces, which are masked by noncontractile tissue at the level of whole musculotendon force. We removed noncontractile tissue force contributions from the recorded musculotendon force using an exponentially-elastic tissue model. We then show that the remaining estimated muscle fiber force resembles history-dependent muscle spindle firing rates recorded simultaneously. These forces also resemble history-dependent forces recorded in stretch of single activated fibers and attributed to muscle cross-bridge mechanisms (Campbell and Moss, 2000). Our results suggest that history-dependent muscle spindle firing in both rats and cats arise from stretch of cross-bridges in muscle fibers.

## Introduction

Muscle spindles are sensory organs within skeletal muscles that are crucial for sensing body segment position and motion (Prochazka and Ellaway, 2012) with mechanosensory signaling characteristics that generalize across species (Vincent et al., 2017). In relaxed muscle, muscle spindle Ia afferents fire when muscles are stretched by an external load, beginning with a high-frequency initial burst of firing, followed by increased firing related to stretch velocity and amplitude in cats, rats, rabbits, and humans (Blum et al., 2017; Proske and Stuart, 1985; Vincent et al., 2017). However, these responses are history dependent, such that the first stretch in a series of identical stretch-shorten cycle elicits an initial burst and response to ramp stretches that are absent or reduced in subsequent stretches (Blum et al., 2017; Haftel et al., 2004; Matthews, 1972; Proske and Stuart, 1985).

In anesthetized cats, history-dependent muscle spindle firing rates mirror history-dependent whole musculotendon forces during muscle stretch. The fine temporal details of Ia afferent firing rates can be precisely reproduced through weighted pseudolinear combinations of recorded whole musculotendon force and dF/dt signals (Blum et al., 2017). As such, elevated whole musculotendon force and dF/dt in the first stretch directly explain the initial bursts and elevated firing rates in the first stretch in a series of stretch-shorten cycles.

In anesthetized rats, however, the history-dependent muscle spindle firing rates cannot be directly explained by history-dependence in whole musculotendon forces (Haftel et al., 2004). Specifically, the force profiles are qualitatively different the muscle spindle firing rates and the forces observed in cat muscle, exhibiting an exponential increase in force during ramp stretches. As longer strain was elicited in prior rat vs cat studies (7% vs. 3%L_o_) greater non-contractile tissues are likely engaged underlying the exponential rise in musculotendon force during stretch (Meyer and Lieber, 2011).

Here we hypothesized that history-dependent muscle fiber forces are masked by noncontractile tissue forces when examining whole musculotendon force during muscle stretch in anesthetized rats. We further hypothesized that muscle fiber forces in rats exhibit similar history-dependence as muscle spindle Ia firing rates recorded simultaneously. We made the simplifying assumption that noncontractile tissue force and muscle fiber force act in parallel, contributing additively to whole musculotendon force. Noncontractile forces were estimated using a simple exponential tissue model, and muscle fiber force was estimated by analytically removing noncontractile tissue force from recorded musculotendon force. We found that a large proportion for whole musculotendon force to be carried by noncontractile tissues. Further, the estimated muscle fiber force and its first time derivative, dF/dt, was history-dependent and closely resembled history-dependent muscle spindle firing rates recorded simultaneously. Our work suggests that the same cross-bridge mechanisms underlie history-dependent muscle spindle firing and muscle fiber forces in rats and cats.

## Methods

### Animal care

All procedures and experiments were approved by the Georgia Institute of Technology’s Institutional Animal Care and Use Committee. Adult female Wistar rats (N=5; 250 - 300 g) were studied in terminal experiments only and were not subject to any other experimental procedures. All animals were housed in clean cages and provided food and water ad libitum in a temperature- and light-controlled environment in Georgia Institute of Technology’s Animal Facility.

### Terminal physiological experiments

Experiments were designed to measure the firing of individual muscle afferents in response to muscle stretch *in vivo* using electrophysiological techniques as documented previously (e.g. (Vincent et al., 2017)). Briefly described, rats were deeply anesthetized (complete absence of withdrawal reflex) by inhalation of isoflurane, initially in an induction chamber (5% in 100% O_2_) and, for the remainder of the experiment, via a tracheal cannula (1.5–2.5% in 100% O_2_).

The triceps-surae muscle group (medial and lateral gastrocnemius and soleus muscles) in the left hindlimb was dissected free of surrounding tissue and detached at its insertion together with a piece calcaneus bone. The severed insertion of the left triceps-surae muscle group and securely attached directly to the lever arm of a force and length-sensing servomotor (Model 305B-LR, Aurora Scientific Inc.), which provided for controlled muscle stretch while recording muscle length and force (dual-mode lever arm system, Aurora Scientific). Initial muscle tension was set to 0.1 N, which is the approximate passive tension observed for ankle and knee angles of 90° and 120°, respectively.

Triceps surae sensory axons were randomly sampled by intra-axonal penetration in dorsal rootlets and selected for detailed study when identified as group Ia based on several criteria (Vincent et al., 2017). First, selected afferents were identified as low threshold mechanoreceptors supplying triceps surae muscles when electrical stimulation of triceps surae nerves generated orthodromic action potentials with conduction delay <2.5 ms. Second, muscle spindle afferents were differentiated by their response to electrically-evoked isometric twitch contractions, which resulted in cessation of stretch-evoked firing in distinction with group Ib afferents which accelerated firing. Finally, group Ia afferents were distinguished from muscle-spindle group II afferents when responded with high-frequency firing (>100 Hz) upon stretch of resting muscle and with high fidelity firing at 100Hz during triceps surae tendon vibration at 100 Hz.

Firing rates and patterns of Ia afferents were studied in response to length-servo controlled stretches applied to triceps surae muscles at rest, i.e. not engaged in active contraction. Triangular stretches, i.e. ramp-release stretch at constant velocity (4 mm/s) were applied with amplitudes ranging from 0.5 to 3 mm. Trials of sequential 3-5 stretches were repeated in between rest periods, i.e. muscles held at resting length for at least 10 seconds allowing expression of history dependent firing responses (Fig 1B; (Haftel et al., 2004)). Whole-muscle force, length changes, and Ia-afferent action potentials were simultaneously recorded at a sampling rate of at least 20kHz and down-sampled to at least 1kHz for analysis. Yank was calculated as the numerical derivative of recorded force. Ia-afferent action potential discriminated for off-line analysis of firing rates and patterns.

**Figure 1:**
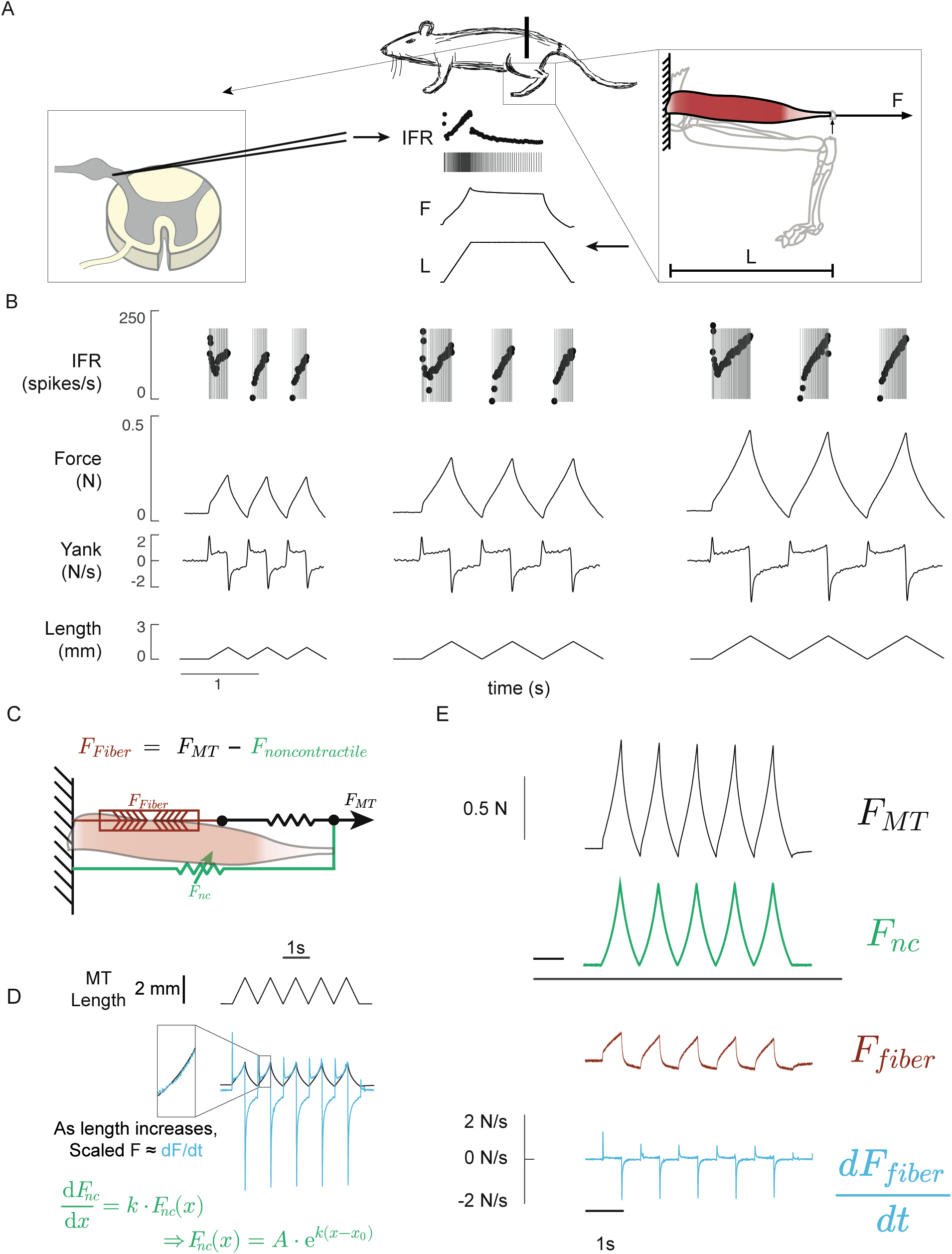
Methods for recording muscle spindle firing rates and estimating muscle fiber force during stretch of relaxed rat muscle. A) Intra-axonal recording from muscle spindle Ia afferents from in dorsal rootlet were recorded during stretch of the triceps surae musculotendon. Muscle length (L) and total musculotendon force (F; panel on right) were recorded with membrane potentials; instantaneous firing rate were computed based on action potential events (IFR; traces in center panel). B) Sets of identical triangular stretches were imposed at three lengths (1 mm, 2 mm, and 3mm). Action potentials (gray lines) with superimposed instantaneous firing rate (black dots) are shown temporally aligned with simultaneously recorded muscle force, yank and length traces. In each trial, muscle spindle Ia firing response to identical stretch stimuli different in the first stretch when compared with subsequent stretches. Specifically, the response to the first stretch contains an initial burst of spikes at stretch onset and more pronounced dynamic response post-initial burst. C) The recorded musculotendon force was assumed to be comprised two additive force components: one component arising from noncontractile tissues (green) with an elastic force-length relationship and another component arising from the muscle fibers (red) and series elastic component (black). D) The recorded musculotendon force (black) and calculated dF/dt (cyan) signals exhibited an exponential force-length relationship which was more dominant at longer stretch lengths, justifying the model of noncontractile tissue as an exponential spring. E) Subtracting the estimated noncontractile force component (green) from the recorded musculotendon force (black) yielded the estimated muscle fiber force (red) and dF/dt (blue), both of which were history dependent, with a larger response during the first stretch.

### Muscle fiber force estimation

To isolate the component of recorded musculotendon force arising from muscle fibers, we assumed noncontractile tissues arranged in parallel muscle fibers (Fig. 1C). We assumed noncontractile tissues were purely elastic with both exponential and linear stiffness (Fig. 1D) describe by the following equation:

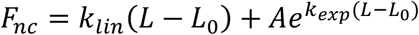

Where L is the entire musculotendon length, L_0_ is the musculotendon resting length, and *K*_*lin*_, *A*, and *K*_*exp*_ spring constants for linear and exponential elements, respectively. Noncontractile tissue forces were then subtracted from the recorded force to estimate the muscle fiber force, which resembled muscle spindle IFRs (Fig. 1E). For each muscle (N = 5), *K*_*lin*_, *A*, and *K_exp_* were each set to values of 0.5 and optimized such that resulting estimated muscle fiber force, when multiplied by a constant, could explain the maximum amount of variance of the recorded Ia afferent IFR during 2mm or 3mm stretch trials (Blum et al., 2017). Parameters from one stretch in each animal was then applied to estimate noncontractile and muscle fiber force in all stretches of that muscle.

### Variance accounted for of musculotendon force by estimated noncontractile force

We estimated the contribution of noncontractile tissues to the total recorded musculotendon force at each perturbation length for a muscle. We used variance accounted for, defined as:

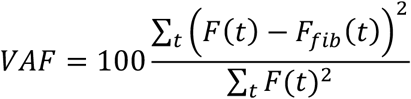

where the numerator represents the sum of squares of the noncontractile tissue model for a given trial and the denominator is the total sum of squares of the recorded musculotendon force for the same trial.

### Estimated driving potential of muscle spindle afferents

To demonstrate that, in principle, arbitrary combinations of estimated muscle fiber force and yank could explain muscle spindle afferent firing rates, we added force and yank together in a pseudolinear manner (Blum et al., 2017):

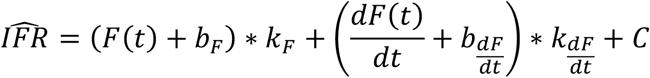

*K_F_* and *K_dF/dt_*, respectively, are constant weights on estimated fiber force and dF/dt and *b_F_* and *b_dF/dt_*, respectively, are constant offsets on fiber force and dF/dt. *C* is a constant offset. For visualization, we chose these parameters to be *K_F_* = 1393 spikes/Ns, *K_dF/dt_* = 3.6 spikes/N, *b_F_* = 10 N, *b_dF/dt_* = 5N/s, and *C* = 34 spikes/s. Additionally, the yank and force components were subjected to a force threshold to emulate the spiking threshold of the neuron.

## Results and Discussion

### Muscle spindle firing rates appeared history-dependent in rats but muscultotendon force did not

As described previously, rat muscle spindle Ia afferents exhibited history-dependent firing rates in response to repeated ramp-release stretch perturbations (Haftel et al., 2004). Initial bursts were observed at the onset of the first stretch and firing during the first stretch was compared to subsequent stretches (Fig 1B). In contrast, simultaneously recorded musculotendon force did not appear history-dependent, apart from a small rise in force at the onset of stretch primarily visible in the yank time series (Fig. 1B) and exhibited an exponentially increasing force with stretch length (Fig. 1C).

### Estimated muscle fiber force exhibited history dependence analogous to that observed in muscle spindle spike rates

Non-contractile force contributions to whole musculotendon force increased with stretch length (Fig 1D). Both the whole musculotendon force and dF/dt signal exhibited a similar nonlinear rise with applied, a property of an exponential relationship (Fig 1D). The non-contractile component was thus modeled by linear and exponential parameters of *K*_*lin*_ = 0.0497 ± 0.0388, *A* = 0.0454 ± 0.317, *K*_*lin*_ = 1.071 ± 0.362 (mean ± sd.; N=5 muscles). Using a single set of parameters for each muscle analyzed, the estimated noncontractile force component accounted for 53 ± 3% (mean ± S.E.M.) of the total variance in force for 3 mm stretches, 34 ± 3% for 2mm stretches, 12 ± 1 % for 1mm stretches, and only 3 % (based on data from one muscle) for 0.5 mm stretches.

The remaining forces represent the estimated muscle fiber force over time, which exhibited history-dependence (Fig. 1E). The estimated muscle fiber force had a pronounced initial force rise and larger initial peak in the first stretch (Fig. 1E). This was accompanied by a larger peak in dF/dt in the first stretch (Fig. 1E). Additionally, this rapid force increased resulting in a higher overall level of force during the first ramp stretch compared to subsequent stretches (Fig 1E).

The history-dependence of muscle spindle firing rates, whole musculotendon force and dF/dt, the and estimated muscle fiber force and dF/dt are illustrated by differences in the first stretch response versus subsequent responses (defined in Fig. 2A). Muscle spindle firing rates were dramatically different on the first stretch compared to subsequent stretches, which were all similar to each other (Fig. 2B, compare yellow to other colored traces). However, total musculotendon force and yank were quite similar across all stretches with only small differences between the first and subsequent stretches (Fig. 2C). Similarly, the estimated muscle fiber force and yank were clearly differentiated on the first ramp-release stretch (Figure 2D, right panel, compare yellow to other colored traces). However, neither muscle fiber force nor yank by themselves resembled the history-dependence of the muscle spindle firing rates.

**Figure 2:**
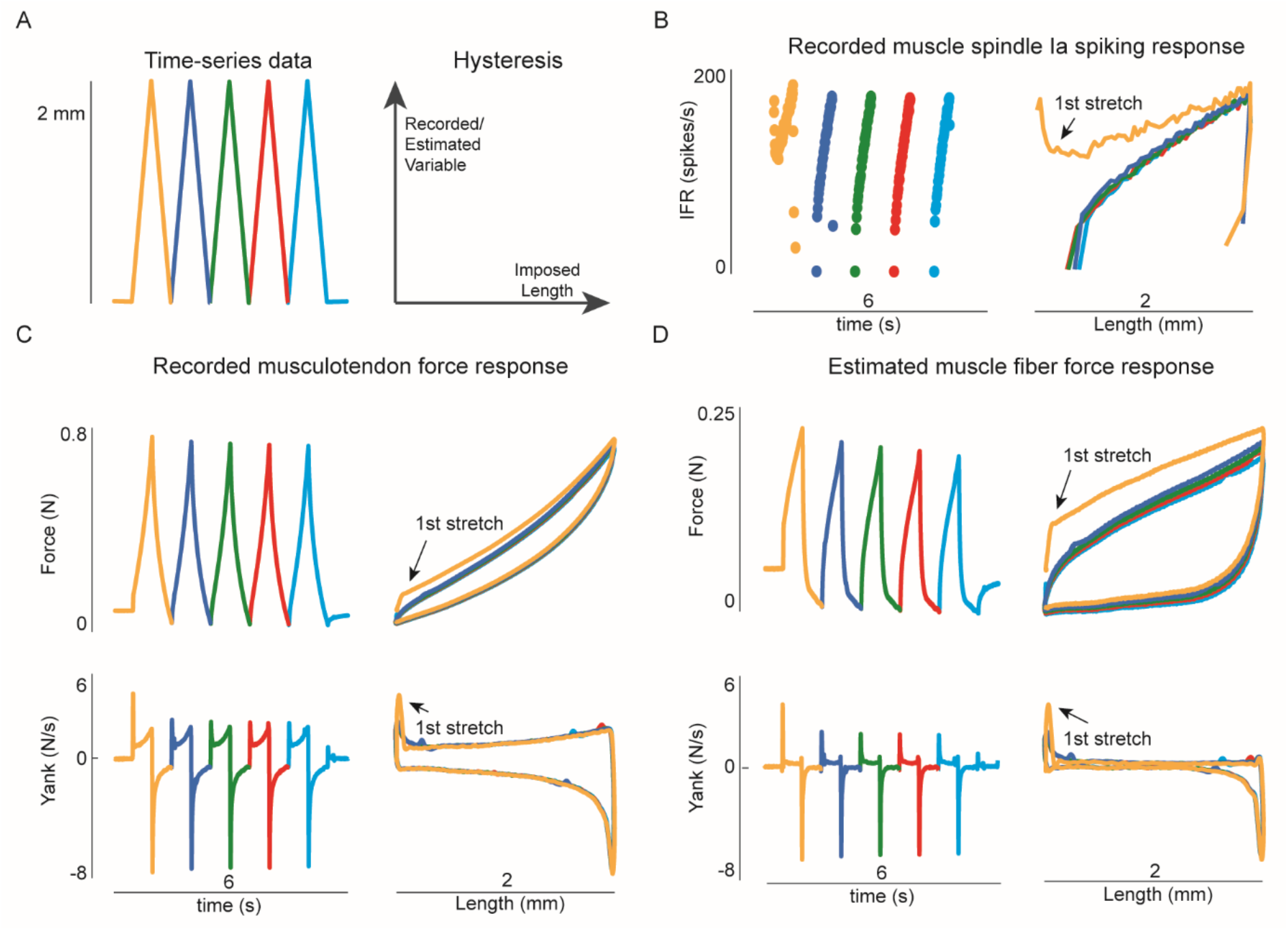
History-dependence of muscle spindle firing rates, musculotendon force and dF/dt, and muscle fiber force and dF/dt. A) Applied length depicted as time series (left panel). Different colors represent different stages of the perturbation. B) Recorded firing rate from Ia afferent to applied length from A. Represented as time series (left) and as a function of length (right). C) Recorded musculotendon force (top) and dF/dt (bottom) D) Estimated muscle fiber force (top) and dF/dt (bottom) estimated from musculotendon force in C.

We found that the history dependence seen in muscle spindle firing rates (Fig. 3A, B) resembled linear combinations of muscle fiber force and dF/dt, subject to a threshold (Fig. 3C, D). Although muscle spindle firing rates in 0.5 mm stretches (e.g. Fig 3B) were qualitatively different than 2 mm stretches (Fig 3A), they were still similar to the linear combinations of the estimated muscle fiber force and yank (Fig. C, D). Morevoer, these estimates were generated using the same weighting of force and yank at both stretch lengths (Fig. 3 E, F). This robustness is all the more remarkable because the same noncontractile tissue properties were used to estimate muscle fiber force and dF/dt (Fig 3 G, H) at different stretch lengths where the amplitude of the noncontractile forces differed dramatically (30.2% vs 3.1% VAF of the total musculotendon force, respectively). In particular the flatter muscle spindle response during the first ramp-release (Fig. B, yellow trace) was also present in the linear combination of muscle fiber force and yank (Fig. 3D, yellow trace).

**Figure 3:**
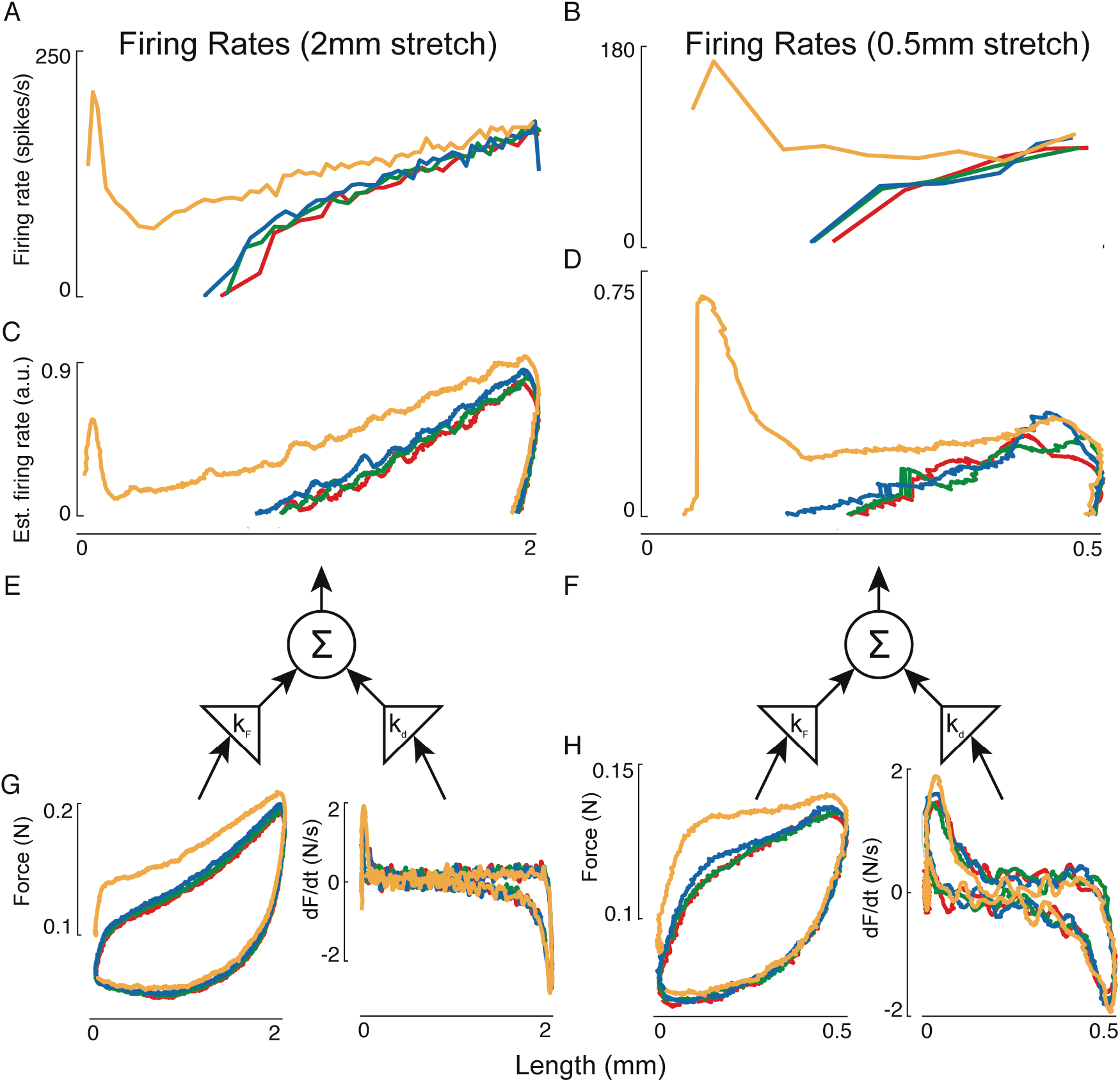
History-dependent muscle spindle firing rates at different applied lengths resembled linear combinations of muscle fiber force and dF/dt. Muscle spindle firing rate for the afferent in A) 2 mm and B) 0.5 mm stretches. C, D)Linear combination of estimated muscle fiber force and dF/dt subject to a threshold exhibited the same qualitative features in both stretch conditions. E,F) The same weights were used in both stretch amplitudes to combine G,H) estimated muscle fiber force and dF/dt signals.

In summary, we show in anesthetized rats that history-dependent muscle fiber forces during stretch of relaxed muscle can be identified from whole musculotendon force even when the majority of that force arises from stretch of non-contractile tissues. Exponential non-contractile force dominated the musculotendon force and yank traces, consistent with reports that muscle fibers carry as little as 15% of total musculotendon loads in rodents ((Meyer and Lieber, 2011), Meyer and Lieber, in review). The remaining residual force was a relatively small component of the total force but resembled characteristics of isolated, activated, muscle fibers in ramp-release stretches (Campbell and Lakie, 1998; Campbell and Moss, 2000; Campbell and Moss, 2002), with an initial short-range stiffness (Getz et al., 1998), as well as higher mean force during the first ramp stretch.

Further, estimated muscle fiber forces exhibited history-dependent characteristics that were similar to history-dependence in muscle spindle afferents recorded simultaneously, suggesting similar muscle cross-bridge mechanisms in both cats and rats. These properties have also been shown to arise from muscle-cross bridge dynamics and only occur in the presence of Ca++ (Campbell and Moss, 2002). The firing rates of muscle spindle primary afferents are directly related to the intrafusal muscle fibers of the spindle, with only a small amount of connective tissue force contributing to the spike-generating force (Boyd et al., 1977; Hunt and Ottoson, 1975; Hunt and Wilkinson, 1980). In relaxed muscle we assume that the intrafusal and extrafusal muscle force exhibit similar responses to stretch, and therefore used our overall estimate of muscle fiber force as a proxy for intrafusal fiber force. As such our work shows that both rat and cat muscle spindles exhibit history-dependence that arises directly from muscle cross-bridge dynamics when stretched, driving the firing behavior.

Our study further provides a neuromuscular explanation for the tendency of human subjects to underestimate forces at longer muscle lengths due to the stretch of non-contractile tissues. Psychophysical studies demonstrate that human participants tend to only perceive forces generated by muscle fibers, and not passive components of force (Tsay et al., 2014). This sense of “fiber-only” force is consistent with the finding in the present study that muscle spindles only fire in response to the force in muscle fibers, but not in non-contractile tissues. While Golgi tendon organs also sense active muscle force, they are located in-series with both muscle fibers and passive connective tissues and are unable to differentiate contractile and non-contractile force. The high contribution of non-contractile forces to whole musculotendon force during muscle stretch in rats could also explain the fact that Golgi tendon organ Ib afferents fire robustly to stretch in rats, but not cats (Vincent et al., 2017).

Taken together, our work suggests muscle spindle firing rate reflect muscle fiber forces during passive stretch across different species. The dissociation of the encoding of contractile and non-contractile forces may be important in understanding the role of proprioceptors in sensorimotor control, and the similarities and differences across species (Vincent et al., 2017). Differences in prior experiments between cats and rats may simply be due to difference in the relative amplitude of stretches, as larger stretches in cat muscle do engage non-contractile force elements that have an exponential characteristic (Matthews, 1931) and would likely need to be considered for larger stretch amplitudes. Indeed, when homologous muscles in the cat, rat, and mouse are stretched with similar strain, firing responses of Ia afferents in each species are similar despite differences in the shape of musculotendon force (Carrasco et al., 2017). Further, in our anesthetized conditions, intrafusal forces within the muscle spindle encoding region is assumed to be similar to the extrafusal muscle force; however, intrafusal muscle force would be increased in conditions where gamma motor neurons innervating intrafusal muscle fibers are active. However, we hypothesize that the fundamental idea that muscle spindles fire in response to intrafusal muscle force and yank will generalize across both passive and active movement conditions.

## References Cited

Blum, K. P., Lamotte D’Incamps, B., Zytnicki, D. and Ting, L. H. (2017). Force encoding in muscle spindles during stretch of passive muscle. PLoS Comput. Biol. 13, e1005767–24.

Boyd, I. A., Gladden, M. H., McWilliam, P. N. and Ward, J. (1977). Control of dynamic and static nuclear bag fibres and nuclearbag fibres and nuclear chain fibres by gamma and beta axons in isolated cat muscle spindles. J. Physiol. (Lond.).

Campbell, K. S. and Lakie, M. (1998). A cross-bridge mechanism can explain the thixotropic short-range elastic component of relaxed frog skeletal muscle. J. Physiol. (Lond.) 510 (Pt 3), 941–962.

Campbell, K. S. and Moss, R. L. (2000). A thixotropic effect in contracting rabbit psoas muscle: prior movement reduces the initial tension response to stretch. J. Physiol. (Lond.) 525 Pt 2, 531–548.

Campbell, K. S. and Moss, R. L. (2002). History-dependent mechanical properties of permeabilized rat soleus muscle fibers. Biophysical Journal 82, 929–943.

Carrasco, D. I., Vincent, J. A. and Cope, T. C. (2017). Distribution of TTX-sensitive voltage-gated sodium channels in primary sensory endings of mammalian muscle spindles. Journal of Neurophysiology 117, 1690–1701.

Getz, E. B., Cooke, R. and Lehman, S. L. (1998). Phase transition in force during ramp stretches of skeletal muscle. Biophysical Journal 75, 2971–2983.

Haftel, V. K., Bichler, E. K., Nichols, T. R., Pinter, M. J. and Cope, T. C. (2004). Movement reduces the dynamic response of muscle spindle afferents and motoneuron synaptic potentials in rat. Journal of Neurophysiology 91, 2164–2171.

Hunt, C. C. and Ottoson, D. (1975). Impulse activity and receptor potential of primary and secondary endings of isolated mammalian muscle spindles. 252, 259–281.

Hunt, C. C. and Wilkinson, R. S. (1980). An analysis of receptor potential and tension of isolated cat muscle spindles in response to sinusoidal stretch. J. Physiol. (Lond.) 302, 241–262.

Matthews, B. H. C. (1931). The response of a muscle spindle during active contraction of a muscle. The Journal of Physiology 72, 153–174.

Matthews, P. B. C. (1972). Mammalian muscle receptors and their central actions.

Meyer, G. A. and Lieber, R. L. (2011). Elucidation of extracellular matrix mechanics from muscle fibers and fiber bundles. J Biomech 44, 771–773.

Meyer, G.A. and Lieber, R.L. (in review).

Nichols, T. R. and Cope, T. C. (2004). Cross-bridge mechanisms underlying the history-dependent properties of muscle spindles and stretch reflexes. Can. J. Physiol. Pharmacol. 82, 569–576.

Prochazka, A. and Ellaway, P. (2012). Sensory systems in the control of movement. Compr Physiol 2, 2615–2627.

Proske, U. and Stuart, G. J. (1985). The initial burst of impulses in responses of toad muscle spindles during stretch. J. Physiol. (Lond.) 368, 1–17.

Tsay, A., Savage, G., Allen, T. J. and Proske, U. (2014). Limb position sense, proprioceptive drift and muscle thixotropy at the human elbow joint. J Physiol 592, 2679–2694.

Vincent, J. A., Gabriel, H. M., Deardorff, A. S., Nardelli, P., Fyffe, R. E. W., Burkholder, T. and Cope, T. C. (2017). Muscle proprioceptors in adult rat: mechanosensory signaling and synapse distribution in spinal cord. Journal of Neurophysiology 118, 2687–2701.

